# Brain Network Analysis in Alzheimer’s Disease and Mild Cognitive Impairment using High-Density Diffuse Optical Tomography

**DOI:** 10.1101/2025.04.28.651132

**Authors:** Emilia Butters, Liam Collins-Jones, Rickson C Mesquita, Deepshikha Acharya, Elizabeth McKiernan, Axel AS Laurell, Audrey Low, Sruthi Srinivasan, John T O’Brien, Li Su, Gemma Bale

## Abstract

Dementia is associated with altered resting state connectivity, measures of which could aid in its early detection and monitoring. High-density diffuse optical tomography (HD-DOT) is well-suited to detect these alterations at scale due to its numerous practical advantages, but it has not yet been applied to dementia. In this study, we investigated resting state functional connectivity across the prefrontal cortex in individuals with Mild Cognitive Impairment (MCI, *n* = 22), Alzheimer’s Disease (AD, *n* = 21), and in healthy controls (*n* = 22). A graph theoretical approach was taken to characterise both global and local patterns of connectivity over a five-minute resting period. We found that individuals with AD and MCI exhibited denser and stronger networks, with shorter path lengths, as well as a loss of consistent key network hubs with worsening cognitive impairment. These results indicate that networks in dementia are more globally connected, albeit in a disorganised manner, perhaps reflecting the recruitment of additional connections for short-term stability which may be ultimately damaging in the long-term. Following the demonstration of HD-DOT’s capacity to detect differences between healthy ageing and dementia, this work opens up new possibilities for the use of optical imaging in the study of this clinical population and HD-DOT’s potential for scalable clinical use.

## 1 Introduction

Dementia is a syndrome which encompasses a range of cognitive symptoms, including problems with memory, executive function, and language, which lead to functional impairment (Arvanitakis et al., 2019). The predominant cause of dementia is *Alzheimer’s Disease* (AD) which is typified by progressive medial temporal lobe (MTL) atrophy and amyloid-*β* and tau deposition throughout the brain. Prior to the marked cognitive impairment characteristic of dementia, individuals experience a period of early cognitive decline known as *Mild Cognitive Impairment* (MCI). This MCI stage is thought to be critical for intervention as it represents a “break point” between the ability to compensate for pathological changes and irreversible functional impairment (Østergaard et al., 2012).

The formal diagnosis of dementia or MCI typically requires scoring below a pre-defined threshold on standardised cognitive tests, indicating a departure from what is considered to be ‘healthy cognitive ageing’. Yet, results on these tests can be highly variable between individuals – they can be influenced by age, education level, and the clinician’s interpretation (Mitchell, 2015) – and how exactly they relate to underlying pathology and disease progression remains unclear. Much effort has been expended to develop a more consistent and objective measure of disease stage primarily through the use of neuroimaging, fluid markers, and genetic testing (Ahmed et al., 2014). In particular, there is considerable focus on the development of biomarkers that rely on tools that are low-cost, accessible, and easily-deployable for widespread clinical use. Such a biomarker, or series of biomarkers, could facilitate early recognition and intervention for those at risk of further cognitive decline and dementia. Early intervention is crucial as disease-modifying drugs appear to be most effective early in the disease course (Van Dyck et al., 2023).

The resting state, or ‘task-free’ condition, is a promising target for biomarker development due to its simplicity and the absence of performance-based confounds. During the resting state, several brain regions exhibit temporally coordinated low-frequency spontaneous neural oscillations referred to as *functional connectivity*. Consistent patterns of these correlations have led to the identification of distinct resting-state networks in the brain (Damoiseaux et al., 2006). Disease-specific alterations in the behaviour of these networks may offer markers for the detection of dementia and provide insights into the early functional changes that may precede later structural changes. Previous studies using functional Magnetic Resonance Imaging (fMRI) have identified altered functional connectivity in AD, primarily in the Default Mode Network (DMN), alongside evidence of a compensatory response characterised by increased connectivity in other networks, such as the ventral attention network (Ibrahim et al., 2021; Li et al., 2014; Vemuri et al., 2012). Similar alterations have been identified in MCI (Binnewijzend et al., 2012) and even in individuals with risk factors for AD but with no cognitive symptoms (Filippini et al., 2009). Additionally, research using Electroencephalography (EEG) has shown a general ‘slowing’ of electrophysiological signals, and reduced signal complexity and synchrony, in AD (e.g. Aoki et al., 2023; Zheng et al., 2023). However, fMRI and EEG cannot be easily nor widely deployed due to their high cost and/or technical demands.

Near-infrared Spectroscopy (NIRS) is an imaging method that detects changes in cortical oxygenation by measuring relative concentration changes of oxygenated (HbO) and deoxygenated haemoglobin (HbR). As NIRS captures the haemodynamic response, it provides an indirect marker of functional brain activation (though it is important to note that these signals can be influenced by underlying vascular dynamics). High-density Diffuse Optical Tomography (HD-DOT) is a method that builds upon the principles of NIRS. In HD-DOT, a high-density array of sources and detectors is used to collect optical data. Unlike traditional NIRS which operates in the channel space, this optical data is then combined with an MRI-derived anatomical head model to resolve three-dimensional maps of cortical oxygenation. These maps can be then be used to quantify functional connectivity between brain regions, just as with fMRI. Previous work has found that connectivity measured by HD-DOT is well-correlated with that measured by fMRI (Eggebrecht et al., 2014) and graph theory metrics have been shown to be reliable in both low-density NIRS (Niu et al., 2012; Novi et al., 2016) and HD-DOT (Uchitel et al., 2022). But unlike fMRI, HD-DOT is low-cost, wearable, silent, portable, easy-to-use, and relatively tolerant of head motion (Pinti et al., 2018). This lack of constraints makes it possible to study the brain outside of the noisy, restrictive environment associated with methods like fMRI, enabling scanning at the bedside, in the home, and with naturalistic experimental paradigms. As HD-DOT also measures two chromophores as opposed to fMRI’s sole measurement of HbR, it consequently offers a promising alternative that combines high spatial and temporal resolution with the portability and scalability needed for widespread clinical use (Butters et al., 2023). HD-DOT is thus an attractive candidate for application to dementia.

To investigate whether HD-DOT can detect alterations in dementia, we examine resting state functional connectivity in individuals with AD, MCI, and in age-matched healthy controls. To our knowledge, this is the first study to apply HD-DOT to these populations. We use graph theory to characterise and quantify patterns of functional connectivity, and then examine the relationship between graph theory-based measures, and severity of cognitive impairment and MTL atrophy. According to previous literature, we expect to observe disrupted functional connectivity in AD with less pronounced network perturbation in MCI. Specifically, we anticipate to observe reduced network efficiency, density, and strength in both clinical groups, with the extent of these changes correlating with the severity of cognitive impairment and MTL atrophy.

## 2 Materials & Methods

### 2.1 Participants

A total of 65 subjects took part in this study. Of these, 21 were diagnosed with AD, 22 were diagnosed with MCI, and 22 were age-matched healthy controls. All subjects in the clinical groups had received a formal diagnosis from a memory clinic and fulfilled the diagnostic criteria for either AD (McKhann et al., 2011) or MCI (Albert et al., 2011). Healthy controls were recruited from the community through flyers, recruitment websites, and the families of study participants. To be included as a control, subjects had to have no history of memory problems and scored above 26 on the Mini-Mental State Examination (MMSE, Folstein et al., 1975). Across all groups, subjects were excluded if they had a condition known, or suspected, to affect cerebral blood flow or haemodynamics. This included individuals with a history of vascular events (e.g. stroke or transient ischaemic attack) or a diagnosed respiratory illness (e.g. asthma or COPD). All subjects underwent a clinical screening interview and neuropsychological testing which included the MMSE and additional scales covering neuropsychiatric symptoms and functional impairment (Table S1). Medial Temporal Atrophy (MTA) was also scored from 0 to 4 for each subject to provide a brain-specific and widely-used measure of disease severity (Scheltens et al., 1992). Scans were scored upon visual inspection by three independent reviewers who had attended a two hour training course but were not expert reviewers e.g. radiologists. The final score was calculated as the sum of the average of their ratings for the left and right hemispheres. The data used in the present work is a subset of that collected as part of the ‘Optical Neuroimaging and Cognition’ study (IRAS ID 319284). This study was approved by Wales Research Ethics Committee. Written informed consent was obtained in person from all subjects, and from informants for subjects in AD or MCI groups.

### 2.2 Experimental paradigm and recording

HD-DOT data was collected during a five-minute resting state period using the LUMO (Gowerlabs Ltd., London, UK). The LUMO is a high-density modular system composed of multi-distance, overlapping NIRS channels. This enables the recording of data from short channels (0-12 mm), which we assume sample the scalp, and long channels (12-40 mm), which we assume sample the cortex. Each module contains three dual-wavelength LED sources (735 and 850 nm) and four photodiode detectors. Data was recorded at 12.5 Hz from 12 modules covering the bilateral frontal cortex for a total coverage of 36 sources and 48 detectors (~1728 possible channels). An appropriately-sized cap (54-56 cm, 56-58 cm, or 58-60 cm) was selected by measuring subjects’ head circumference prior to fitting.

For the majority of subjects (67.1%), data collection took place in their homes. For the remainder of subjects, data collection took place in a clinic room at the University of Cambridge. Although testing locations varied, all recordings were carried out in quiet, dimly-lit environments. During the recording, subjects were seated in a chair and instructed to rest quietly with their eyes closed while remaining awake, minimising head movements beyond small adjustments, and avoiding structured thoughts or tasks. A comparison of data quality, defined as the percentage of good channels per source-detector distance range, revealed no significant differences between the home and clinic settings (Figure A1). During acquisition, a software error in the data-saving process resulted in three datasets being corrupted. These were subsequently re-recorded.

### 2.3 Data pre-processing

The optical data was pre-processed using the Homer2 toolbox (Huppert et al., 2009) in Matlab v2021b (The MathsWorks Inc.; MA, USA). Channels were removed if (1) their mean coefficient of variation was above 8.3% (equivalent to a signal-to-noise ratio *<* 12), (2) their mean signal intensity *>* 1×10 ^11^ V, (3) their source-detector separation *>* 100 mm, as per (Uchitel et al., 2022), or (4) heart rate could not be detected (i.e. if the maximum of the Fast Fourier Transform (Cooley et al., 1969) of that channel was not in the 0.5-2 Hz range). The data was visually inspected for motion artefacts and motion correction was not deemed necessary due to the low motion burden. Raw intensity signals were first converted to optical densities (Δ*OD*) using the Homer2 toolbox (Huppert et al., 2009). The Δ*OD* data was then filtered using a third-order Butterworth bandpass filter (0.01–0.1Hz). Potential contamination of the data by scalp haemodynamics was minimised by regressing the data from the nearest short channel (between 0–12mm) from the long channels. This was done using the DOTHUB toolbox (https://github.com/DOT-HUB/DOT-HUB toolbox).

### 2.4 Source localisation

As head size and shape vary across the population, we cannot assume that the same source-detector pair will sample exactly the same brain region in different subjects. To account for this, photogrammetry was used to digitise the locations of the optodes and cranial landmarks for each subject (as per Vidal-Rosas et al., 2021). This method involves aligning the features from multiple photographs to create a point cloud from which the coordinates of each desired point can then be extracted. To facilitate this, bright green equilateral triangles (~18 mm length per side) were placed on each LUMO module such that the vertex of each triangle overlay a source (Figure 1d). Similarly, bright blue circular stickers (8mm in diameter) were placed over five cranial landmarks: the left pre-auricular point (Al), the right pre-auricular point (Ar), the nasion (Nz), the inion (Iz), and the vertex (Cz), measured as halfway between Nz and Iz. Each point cloud was generated from two 360^°^ videos of subjects’ heads using Agisoft Metashape (Agisoft LLC, St. Petersburg, Russia). The resulting models were then scaled using MeshLab 2023 (Cignoni et al., 2008) according to known dimensions between sources provided by the manufacturer. The approximate coordinates of each source and detector were manually extracted using a custom script written in Matlab v2021b (Figure 1e).

**Figure 1:**
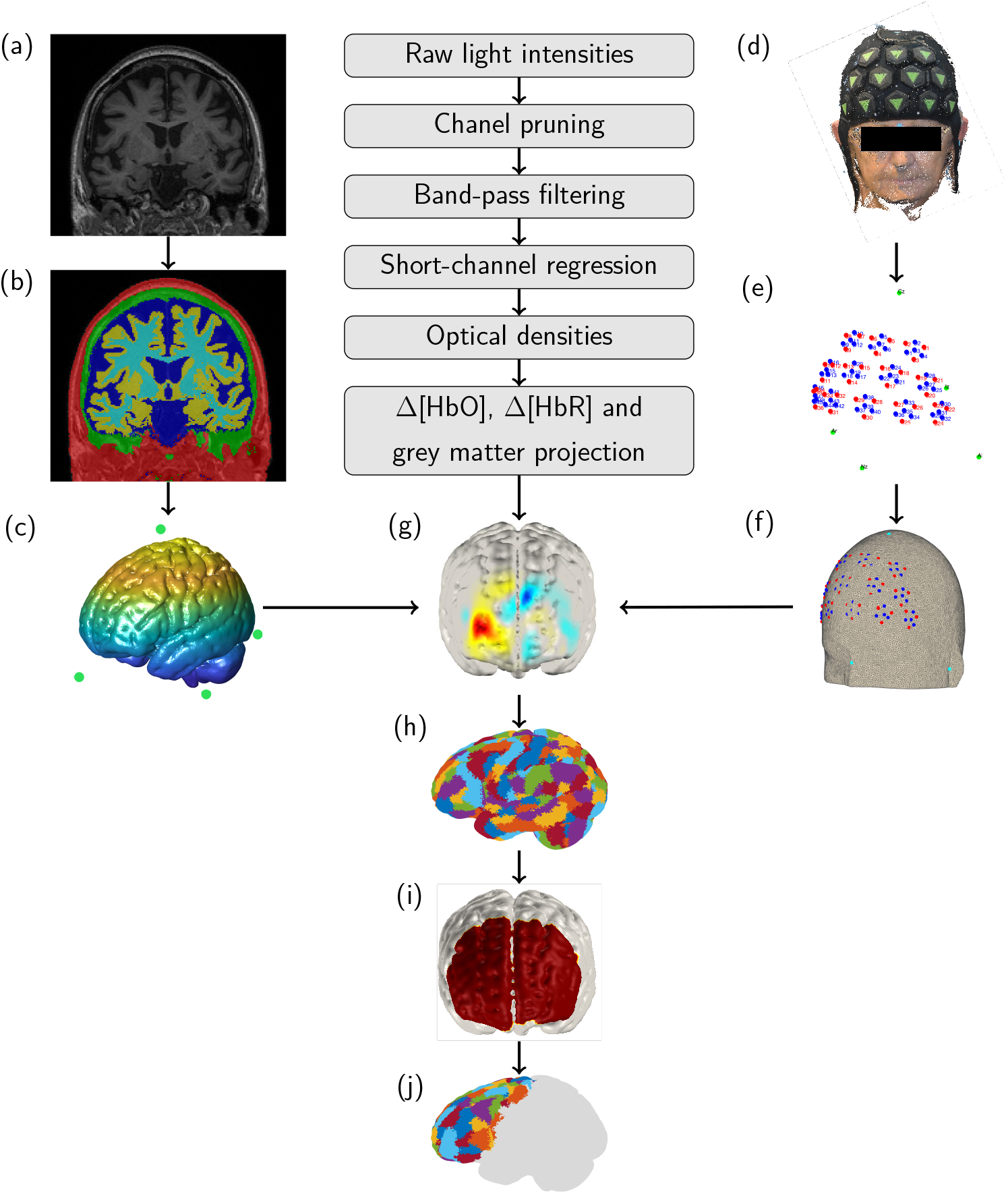
Processing pipeline for the HD-DOT data. (a) An example structural T1-weighted image acquired for subjects in clinical groups. The MNI atlas was used for healthy controls. (b) Segmentation of the structural image into five tissue types using SPM12: scalp, skull, cerebrospinal fluid, white matter, and grey matter. (c) MRI-derived tetrahedral grey matter mesh and cranial landmarks. (d) An example point cloud of a subject wearing the LUMO cap, created using photogrammetry. (e) Manually-extracted optode and cranial landmark coordinates. (f) Optode and cranial landmark coordinates registered to a subject’s tetrahedral mesh. (g) An example topographic reconstruction of cortical oxygenation. (h) Schaefer parcellation atlas registered to native space via non-linear transformation. (i) Sensitivity mask including only nodes sensitive to *>* 5% of the normalised Jacobian. (j) Only parcels with *>* 50% of sensitive nodes included per subject. Time series of each included node averaged per parcel.

### 2.5 Head modelling and registration

To create tomographic maps of brain oxygenation, the high-density NIRS data was reconstructed using a head model. For healthy controls, the MNI-152 template (Maintz & Viergever, 1998) was used, as we assume that these subjects have a typical head size and shape for which standard atlases are sufficient (Ferradal et al., 2013). However, due to the cortical atrophy that is associated with dementia (Mouton et al., 1998), subject-specific MRI-derived head models were used for subjects with AD and MCI. As light propagates through different tissue types in distinct ways, using a subject-specific head model to capture the precise geometry of each tissue type enables the light propagation model to account for pathology-related changes in brain size and shape. This ensures that changes in the optical properties of extra-cerebral tissues are not misattributed to functional changes occurring in the cortex. To create these head models, structural T1-weighted MRI scans were acquired for each subject with AD or MCI (Figure 1a, for acquisition details see Table A1).

Subject-specific head models were created by first segmenting structural MRI using SPM12 (www.fil.ion.ucl.ac.uk/spm) into five tissue types (Figure 1b): grey matter, white matter, cerebrospinal fluid, skull, and scalp. The segmentations were used to generate voxelised tissue masks for each tissue type which were then combined to create a single labelled map per subject. MRI-derived cranial landmarks were manually extracted using ITK-SNAP (Yushkevich et al., 2006), where Cz was estimated. The labelled tissue map and landmarks were then used to construct a three-dimensional tetrahedral mesh using Iso2mesh (Fang & Boas, 2009) in Matlab v2021b (Figure 1c). The optode positions were then registered to the subject’s native space using an affine transformation between the MRI- and photogrammetry-derived cranial landmarks (Figure 1f).

### 2.6 Image reconstruction and parcellation

To reconstruct the optical data, a forward model of light propagation was calculated using the diffusion-approximation to the radiative transfer equation (Arridge, 1999). The sensitivity of the optical signals (*S*) to changes in absorption coefficients was then estimated by deriving a Jacobian matrix (*J*) for each wavelength via the finite element method (Schweiger et al., 1993). This was implemented using Toast++ (Schweiger & Arridge, 2014). To resolve for changes in absorption coefficient, the Jacobian was pseudoinverted via the Moore-Penrose method. A zeroth-order Tikhonov regularisation was applied using a hyperparameter of 0.01 to stabilise the solution. The resulting images were then converted to images of per-node changes in concentration of HbO and HbR (Δ *C*) using *S* = *J* · Δ *C*. A node in the grey matter mesh was defined as sensitive, and thereafter included, if its sensitivity in the Jacobian matrix exceeded 5% of the maximum value of the normalised Jacobian for both wavelengths (Figure 1i, Uchitel et al., 2022). The forward model, its inversion, and image reconstruction were implemented using the DOTHUB toolbox (https://github.com/DOT-HUB/DOT-HUBtoolbox).

To facilitate region-specific analyses and reduce data dimensionality, the reconstructed images were parcellated using the 400 parcel Schaefer parcellation atlas (Figure 1h, Schaefer et al., 2017). This granularity was selected based on preliminary testing, as it effectively reduced data size whilst providing stable and sufficient parcel coverage across subjects for comparing network architectures. In the Schaefer atlas, each node in the grey matter mesh is assigned a functionally-relevant cortical *parcel* (Uchitel et al., 2022). Each parcel in this atlas is also allocated to a resting state network (Yeo et al., 2011). For data reconstructed using a subject-specific head model, the atlas was non-linearly transformed to each subject’s native space using antsRegistration from Advanced Normalisation Tools (Avants et al., 2010) as the atlas is registered to MNI space. Nodes in the grey matter mesh were matched to the closest node in the parcellation atlas using a k-nearest neighbour algorithm. For each subject, a parcel was only included if *>* 50% of nodes in that parcel were sensitive. A single time series for each parcel was then obtained by averaging the time series of all sensitive nodes within that parcel (Figure 1j). Only Δ*HbO* was analysed in the present study as it generally has a higher signal-to-noise ratio and statistical power than Δ*HbR* (Tachtsidis I and Scholkmann F., 2016). Whether the number of sensitive parcels differed per group was determined using an Analysis of Variance (ANOVA, Figure A2).

### 2.7 Graph theory analysis

Functional connectivity was assessed using a graph theory approach. Networks can be represented by graphs: a set of vertices (*V*) connected by edges (*E*). In the present case, the vertices represent brain regions, i.e. parcels, and the edges represent functional connectivity, i.e. time-dependent coordinated activity between these regions. For each subject, functional connectivity was calculated by computing the Pearson’s correlation coefficient (*r*) between the time series of each pair of parcels across the entire five-minute time course (Figure 2b-c). Accordingly, each connectivity matrix had the size *N* × *N* where *N* is the number of sensitive parcels for that subject.

**Figure 2:**
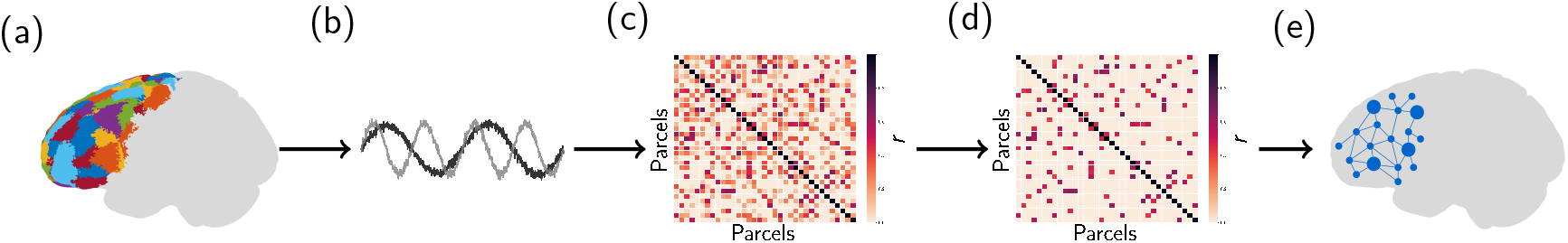
Functional connectivity analysis pipeline. (a) Parcellation of the reconstructed image. (b) Time course extracted from each parcel. (c) Pearson’s *r* correlation matrix calculated for all parcels. (d) Matrix thresholded to remove spurious connections. (e) Graph construction.

For graph analysis, a weighted undirected adjacency matrix was created from the pairwise Pearsons’s correlation between parcels. To do so, correlation matrices was first transformed using Fisher’s r-to-z transform to stabilise variance, then thresholded using an absolute threshold of 0.2 to remove spurious correlations (Novi et al., 2016, Figure 2d). Edge weights were only retained for edges exceeding this threshold and were assigned their z-transformed values. The resulting adjacency matrices were min-max normalised and treated as subject-specific graphs for subsequent analyses.

Following graph construction for each subject (Figure 2e), various metrics were calculated to characterise the connectivity patterns of each graph using the Brain Connectivity Toolbox (Rubinov & Sporns, 2010). Connectivity was assessed both globally and locally. Global network connectivity was evaluated across the entire graph for each subject by averaging graph theory metrics across all of a subject’s sensitive parcels (Figure 3). Connectivity was assessed via degree and centrality (betweenness centrality and Eigenvector centrality), segregation was assessed via clustering coefficient, and flow was assessed via efficiency. Total strength was calculated as the sum of the weights of all edges between nodes in the graph.

**Figure 3:**
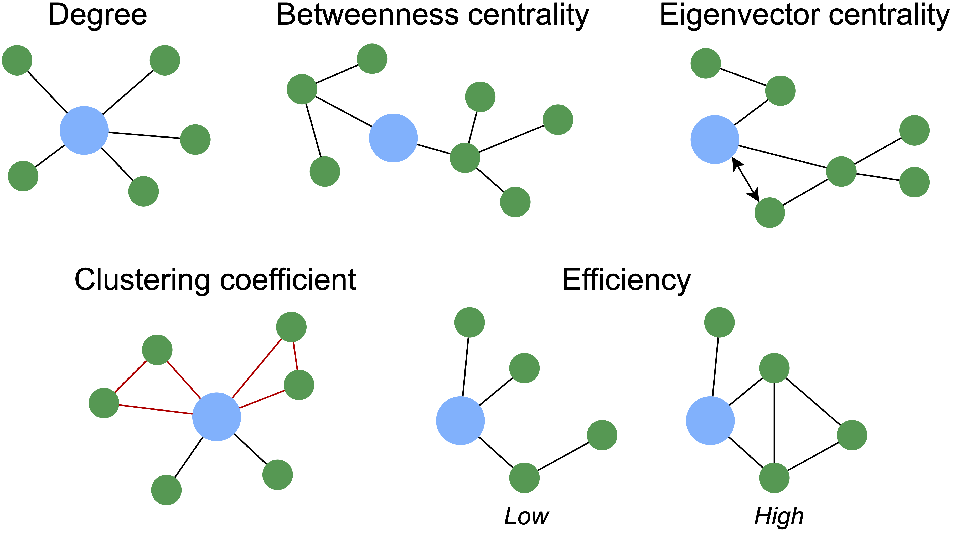
Graph theory metrics. Degree – the number of connections a node has. Betweenness centrality – how important a node is for connecting other nodes. Eigenvector centrality – how well-connected a node’s neighbours are. Clustering coefficient – how likely a node’s neighbours are to be connected to each other. Efficiency – how easily information can flow through a network.

Local connectivity was then assessed by concatenating all subjects’ data parcel-wise to examine regional connectivity patterns in isolation. This means that connectivity was evaluated independently for each parcel, without considering the rest of the network. Clustering properties were first evaluated (Figure 4). Modularity was calculated using the Louvain algorithm (Blondel et al., 2008) which quantifies the strength of community structure in a network by partitioning it into distinct clusters. Following this, the participation coefficient was computed for each parcel as the proportion of its degree that connects it to modules other than the one it belongs to. Participation coefficients were then averaged per resting state network (Yeo et al., 2011), excluding networks present in *<* 20% of subjects (i.e.dorsal attention, visual, somatomotor, and temporoparietal).

**Figure 4:**
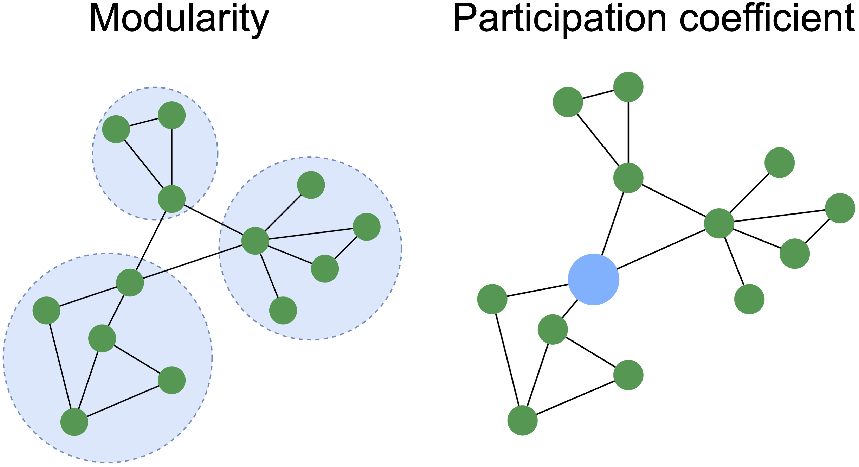
Clustering properties. Modularity – how well a network can be divided into groups. Participation coefficient – how connected a node is to different modules.

**Figure 5:**
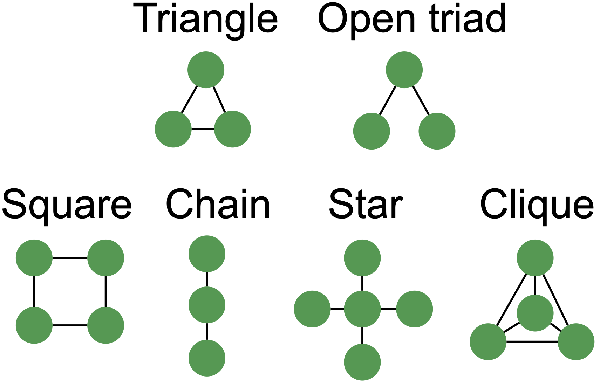
Three-node (top) and four-node motifs (bottom) were considered.

The number of distinct sub-patterns, termed *motifs*, was calculated per graph and subsequently normalised by the total possible number of each motif per graph. Motifs are recurring sub-patterns within a larger network that may reflect functional regularities. The frequency of three-node motifs: triangle and open-triad, and four-node motifs: square, chain, star, and clique, was calculated.

Finally, to identify important network hubs across groups, the frequency with which parcels were found in the top 10% for participation coefficient, degree, and betweenness, and Eigenvector centrality was calculated. An aggregate ‘overall centrality’ score was defined as the median of the three. Parcels with low counts (i.e. those identified in *<* 15% of subjects in each group) were removed and only parcels defined as hubs in at least 20% of subjects were thereafter included. Functional connectomes were created using the NetworkX module (Hagberg et al., 2008) in Python v3.8.8. To do so, thresholded adjacency matrices were min-max normalised across all groups and only significant connections (*p <* .05) present in *>* 20% of subjects were included.

### 2.8 Statistical analysis

Group-level differences in graph theory metrics, demographic variables, and clinical data were calculated following the removal of missing values (if necessary) and assessment of normality using the Shapiro-Wilk test (Shapiro & Wilk, 1965). Depending on the distribution of the data, either parametric or non-parametric statistical tests were used for all group-level and post hoc analyses. Correlations between global graph theory metrics and age, MMSE score, and MTA rating were determined using a Pearson’s or Spearman’s correlation coefficient. Group differences in sex ratio, and the proportion of subjects with a family history of dementia or stroke were assessed using the chi-squared (*χ*^2^) test. Statistical analyses were conducted using either Python v3.8.8, or Matlab v2021b with significance thresholds set at *p <* .05. The Benjamini-Hochberg procedure was used to control the false discovery rate (FDR, Benjamini and Hochberg, 1995) unless otherwise stated.

## 3 Results

### 3.1 Study population summary

A summary of subject characteristics is shown in Table 1. There were no significant differences in age or sex ratio across groups. Full neuropsychological test scores are detailed in Table S1.

**Table 1:** Summary of demographic and clinical data across study groups. Shown as mean ± standard deviation.

| | HC ($n=22$) | MCI ($n=22$) | AD ($n=21$) | p |
| --- | --- | --- | --- | --- |
| Age (y) | 74.5 ± 6.16 | 75.0 ± 6.64 | 72.6 ± 8.42 | 0.40 |
| Female (%) | 50.0 | 45.5 | 42.9 | 0.89 |
| Education (y) | 20.0 ± 3.67 | 18.7 ± 3.31 | 17.8 ± 2.93 | 0.18 |
| Family history of dementia (%) | 59.1 | 68.2 | 66.7 | 0.80 |
| Family history of stroke (%) | 27.3 | 22.7 | 28.6 | 0.90 |
| MTA rating | n/a* | 3.36 ± 1.8 | 4.90 ± 1.50 | **0.01** |
| MMSE score | 28.8 ± 1.15 | 26.5 ± 2.23 | 21.3 ± 4.35 | **0.00**<sup>a,b,c</sup> |
| MoCA | 26.1 ± 2.07 | 23.2 ± 3.45 | 16.3 ± 5.23 | **0.00**<sup>a,b,c</sup> |

P-value indicates overall group-level comparison. Superscript letters denote significant pairwise group comparisons: <sup>a</sup> HC vs MCI, <sup>b</sup> HC vs AD, <sup>c</sup> MCI vs AD. \* MRIs were not acquired for healthy controls so MTA was not rated. MTA rating shown as the sum of left and right hemispheres.

### 3.2 Functional connectivity

#### 3.2.1 Global dynamics

The functional connectomes for each group are shown in Figure 6. A significantly greater proportion of connections survived the threshold of 0.2 in AD and MCI (*p <* .05) compared to in healthy controls (Tukey’s HSD test). Significant differences in global functional connectivity were found across several graph theory metrics between healthy controls and both clinical groups, however no differences were found between MCI and AD groups. Specifically, total strength was higher in AD (*U* = 337, *p <*.05, Figure 7a) and MCI (*U* = 126, *p <* .05) compared to healthy controls, as was average degree density (*t*(2) = 3.06 for AD, *t*(2) = −3.30 for MCI, *p <* .01, Figure 7c) and global efficiency (*U* = 362 for AD, *U* = 94 for MCI, *p <* .01, Figure 7e). Although several of the graph theory metrics scale with the number of parcels present (which may introduce bias between groups), no significant differences were found between the number of sensitive parcels across groups (*H*(2) = 2.10, *p* = 0.35, Figure A2a). In sum, these results indicate *increased* connectivity in AD and MCI, characterised by denser networks, shorter pathlengths, and higher total strength, with no differences between MCI and AD or across measures of network segregation.

**Figure 6:**
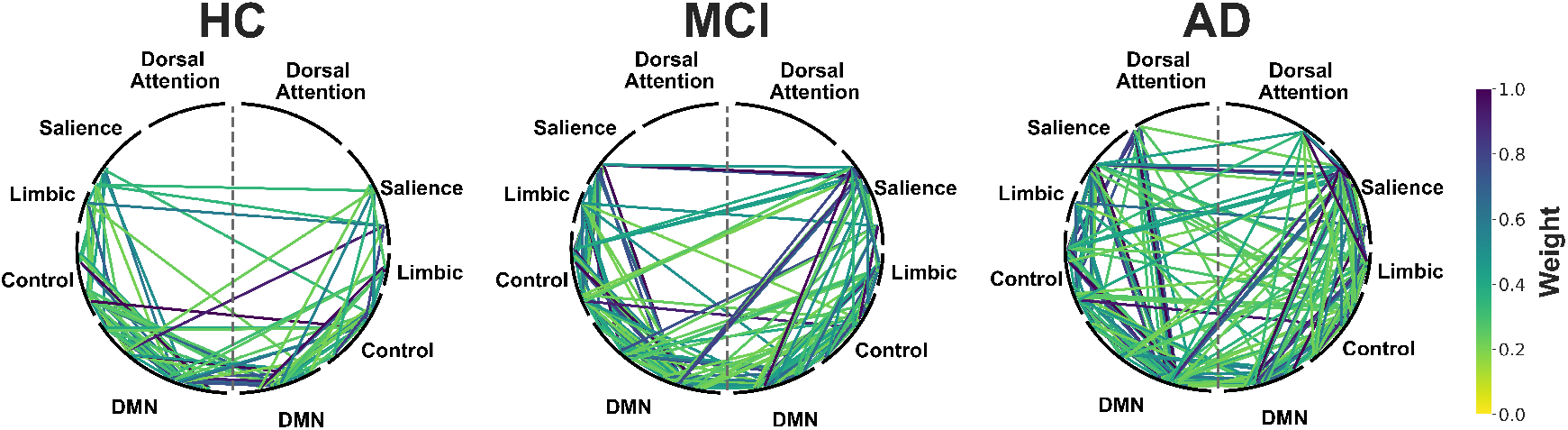
Functional connectomes showing significant connections (*p <* .*05*) present in healthy controls, Mild Cognitive Impairment, and Alzheimer’s Disease. Shown as weighted edges min-max normalised across all groups. Thresholded to only include connections present in *>* 20% of subjects and spurious connections (*r <* 0.2) removed. Parcels grouped according to resting state network (Yeo et al., 2011). Grey dashed line divides left and right hemispheres. DMN = Default Mode Network. *n* = 65.

**Figure 7:**
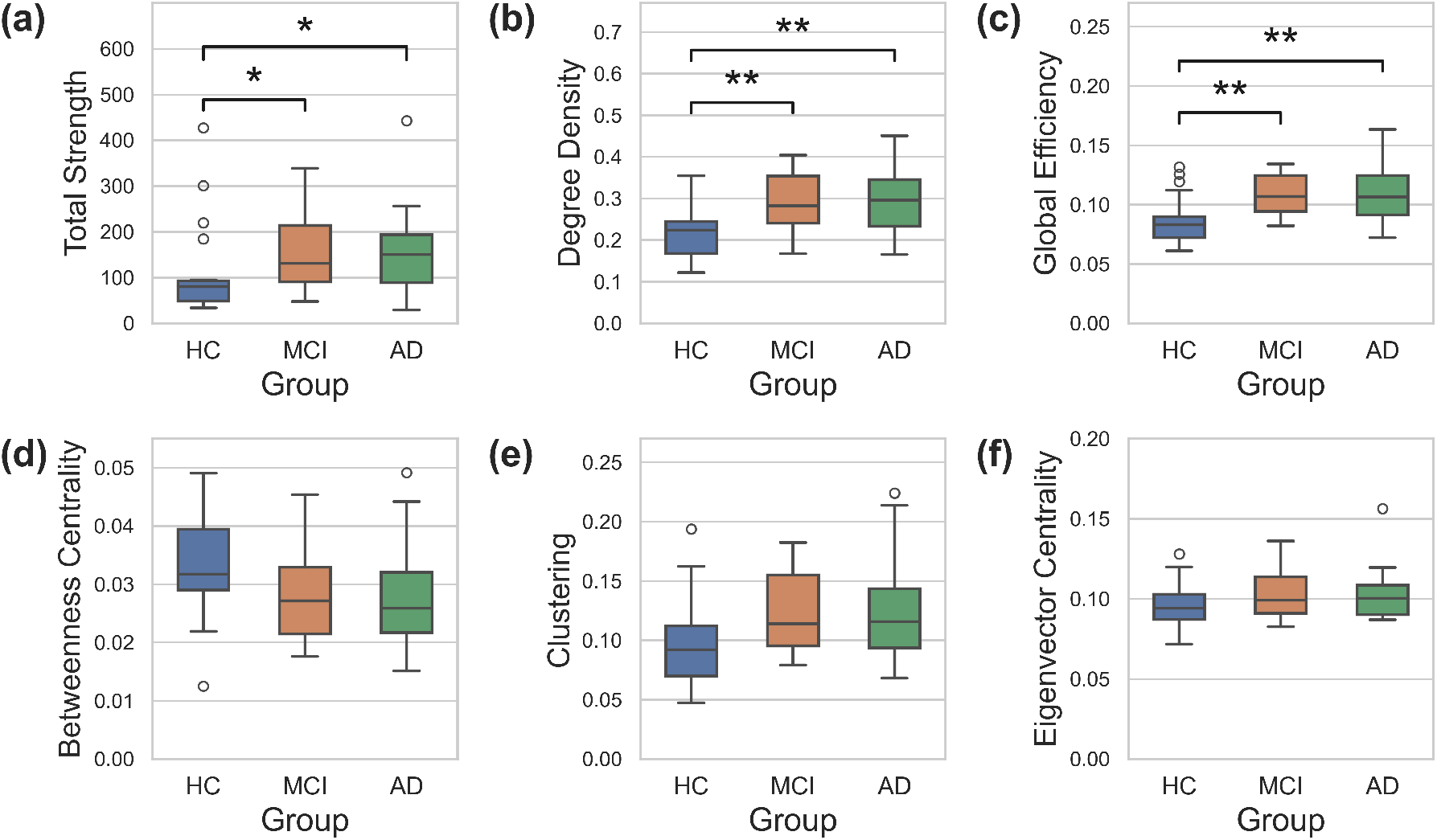
Group-level global functional connectivity, reported as the median with interquartile range. Statistical significance demonstrated by *, *p <* 0.05; **, *p <* 0.01; FDR corrected.

#### 3.2.2 Local dynamics

To identify specific networks which are highly connected, graph theory metrics were also calculated at the local level. Clustering properties were first considered. Modularity did not differ between groups (*H*(2) = 3.78, *p* = 0.15), however, a two-way ANOVA found a main effect of group (Figure 8,*F* (2,236) = 6.05, *p <* .01, *η*^2^ = 0.044) and resting state network (*F* (3,236) = 0.032, *p <* .001, *η*^2^ = 0.088) on participation coefficient. No interaction between group and network was observed (*F* (6,236) = 0.57, *p* = 0.75). Post hoc comparisons revealed overall significantly higher coefficients in AD compared to healthy controls (*p <* .01). This indicates a greater connectivity of parcels with those in clusters outside of their own, as is reflected in the functional connectomes of the AD group (Figure 6). Additionally, lower coefficients in the DMN compared to the control, limbic, and salience networks (*p <* .01) were found across all groups.

**Figure 8:**
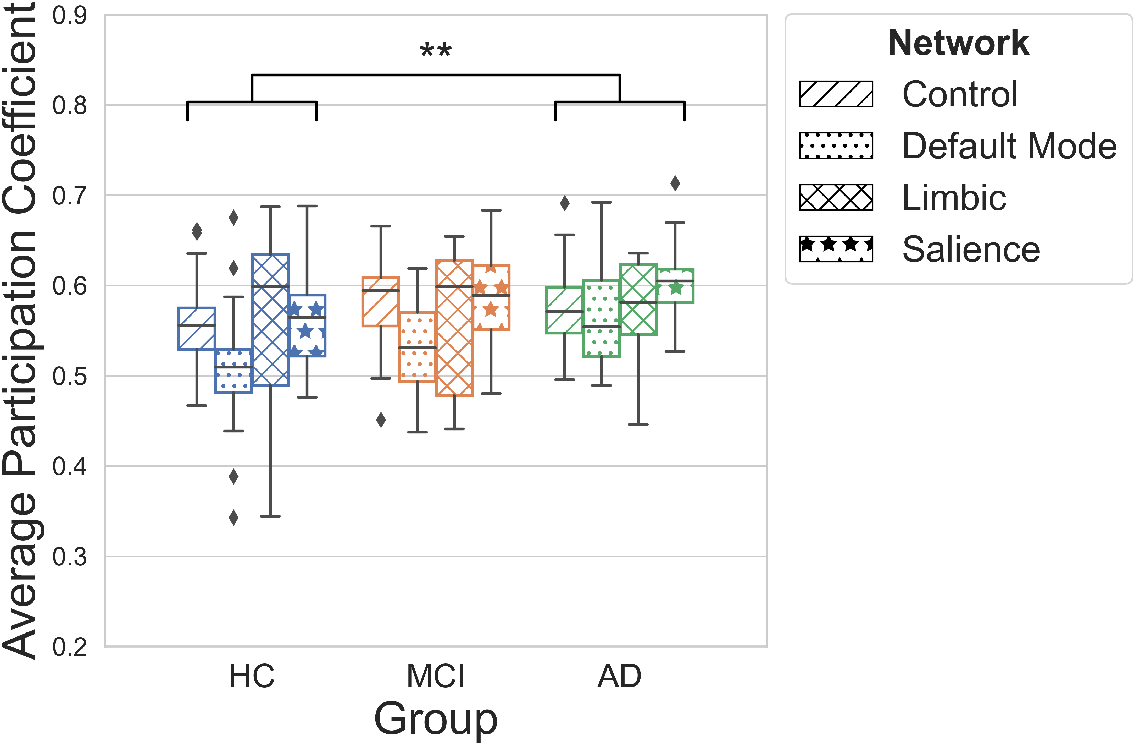
Average participation coefficient per network across groups, reported as the median with interquartile range. A two-way ANOVA found a main effect of group (*p <* .01) and network (*p <* .001). Post hoc comparisons conducted using Tukey’s HSD test. Network differences not shown for visual clarity. *n* = 65.

A graph can also exhibit various motifs, i.e. distinct sub patterns of connectivity between nodes, which can provide insights into a graph’s functional organisation. The prevalence of different types of motifs was quantified and compared across groups (Figure 9). An Aligned-Rank Transformation (ART) ANOVA revealed a main effect of group (*F* (2,434) = 22.88, *p <* .001) and motif type (*F* (2,434) = 257.18, *p <* .001) on the number of motifs present. No interaction between group and motif type was found (*F* (2,434) = 1.63, *p* = 0.08). Post hoc comparisons revealed a significantly higher prevalence of motifs in the AD and MCI groups compared to healthy controls (*p <* .001). As this finding may simply reflect the higher degree density found in AD and MCI, the ratio of triangles (indicating local clustering) to open triads (reflecting a less cohesive architecture) was calculated, however, no difference was found between groups (*H*(2) = 4.70,*p* = 0.1). Taken together, these results mirror the global network changes observed in MCI and AD, with higher participation coefficients in AD, and higher motif counts reflecting increased degree density in clinical groups.

**Figure 9:**
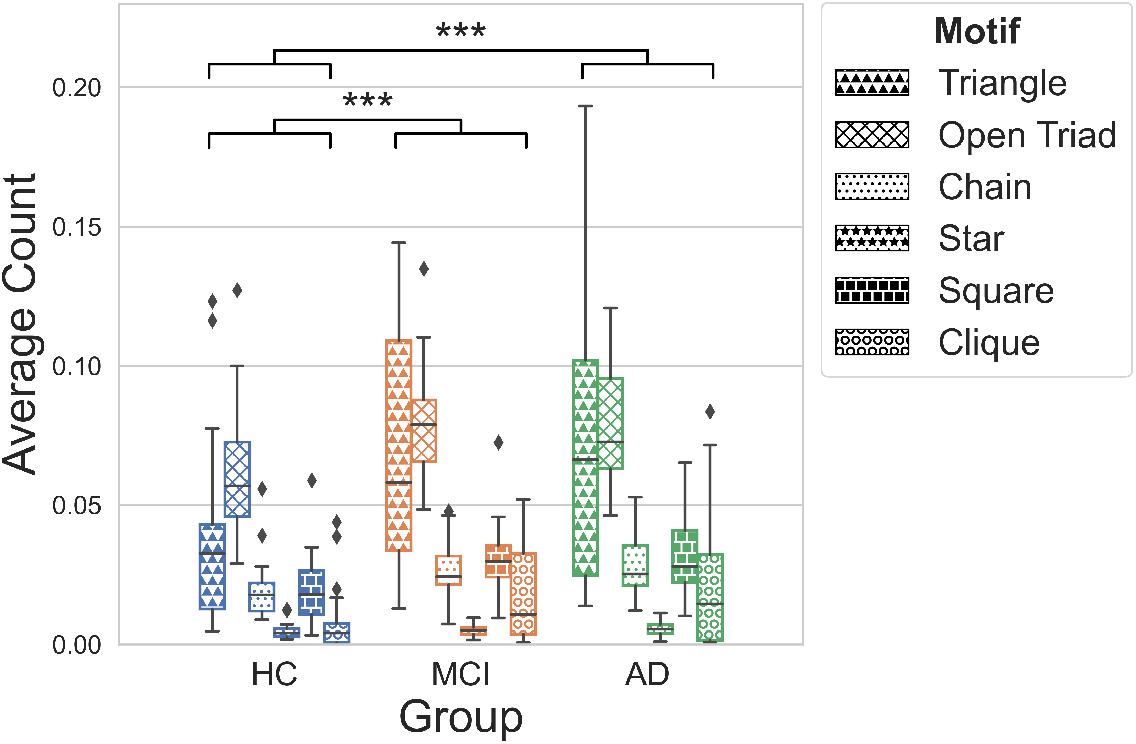
Average count of motif type across groups, reported as the median with interquartile range. An ART-ANOVA found a main effect of group (*p <* .001) and motif type (*p <* .001) on motif count. Post hoc comparisons conducted using Tukey’s HSD test. Motif differences not shown for visual clarity. *n* = 65.

To identify key network hubs within each group, regions exhibiting consistently high functional connectivity across subjects were identified (Figure 10). These hubs generally belonged to the control network and DMN. In healthy controls, parcels with high participation coefficients occurred most frequently in the right hemisphere (72.1%). The opposite was true for MCI and AD, where high-coefficient parcels were primarily found in the left hemisphere (34.3% and 44.2% respectively). Focusing on regions with consistently high centrality measures, the hub which most frequently exhibited high overall centrality across all groups was observed in healthy controls. This was the left lateral PFC, which is part of the control network. Only this region scored highly for all of clustering, efficiency, and centrality across groups, suggesting it plays an influential role in healthy controls but not in clinical groups. The right dorsolateral PFC also showed consistently high degree centrality in healthy controls, and the left dorsal PFC, betweenness centrality. In contrast, the right orbitofrontal cortex within the limbic system was identified as a key region in MCI, exhibiting consistently high overall centrality. In AD, this was the case only for the left medial PFC (part of the DMN), also a key region in MCI but to a lesser extent.

**Figure 10:**
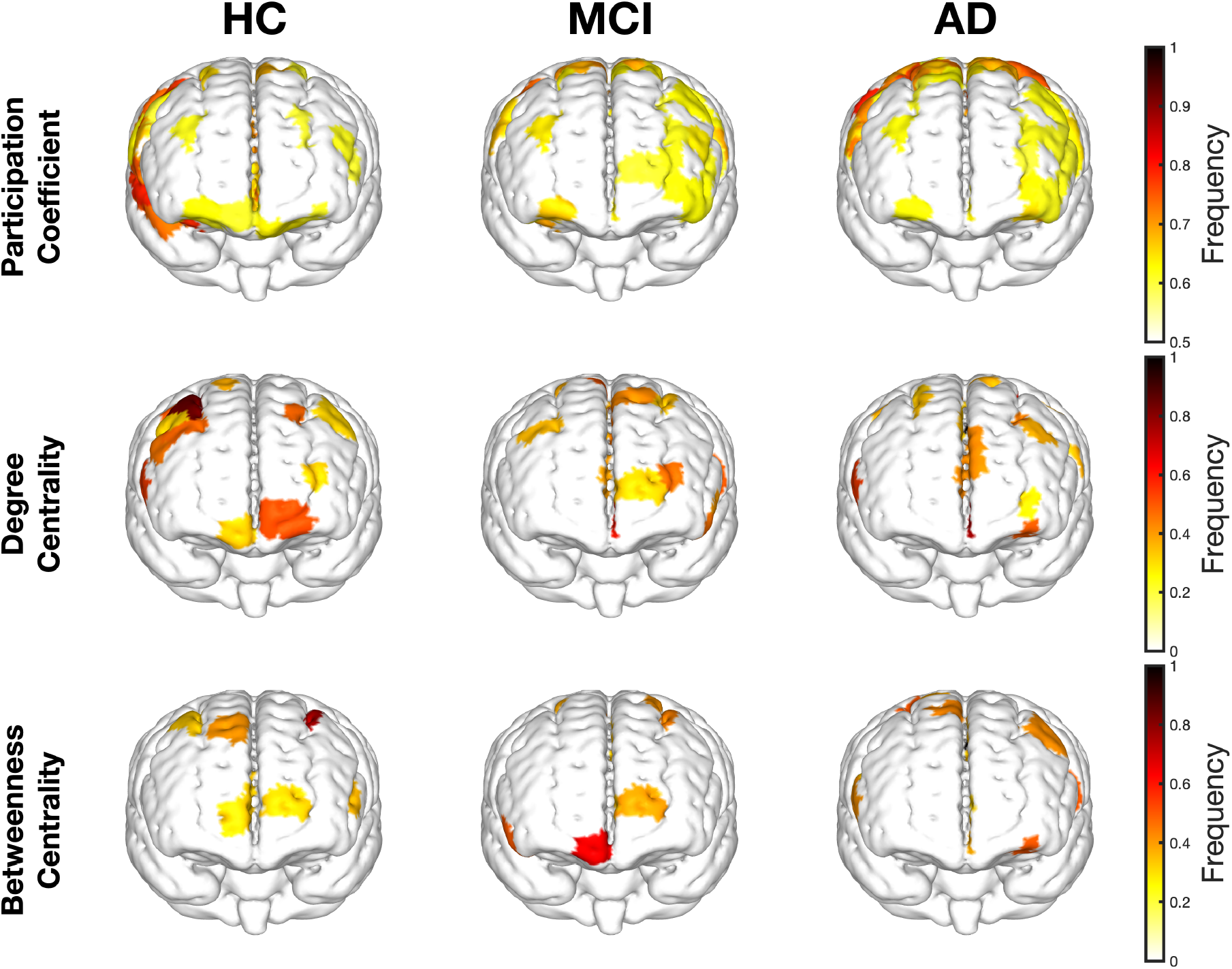
Normalised frequencies of high-scoring parcels per metric. Presented on a standard MNI atlas. Parcels defined as ‘high’ scoring if present in the top 10% for each measure in at least 20% of subjects. Frequency of participation coefficients thresholded to only include frequencies *>* 0.6 for visual clarity. *n* = 65.

Interestingly, a greater number of hubs in the DMN were found in AD and MCI compared to healthy controls for both degree (7:9:4) and betweenness centrality (5:3:4). Only the ventral and lateral PFC, in the DMN and control network respectively, were important hubs for efficiency and clustering in healthy controls and MCI. By contrast, no key networks hubs were identified for clustering or efficiency in AD.

#### 3.2.3 Association with clinical data

There was no association between any of the global metrics and either MTA rating or age (*p >* 0.05) when considering all groups together. Weak to moderate correlations were observed between average betweenness centrality (*r* = −0.36, *p <* .05) and total strength (*r* = −0.35, *p <* .05), and MMSE score (Figure 11). Further weak correlations were found between average degree density (*r* = −0.27, *p <* .05), global efficiency (*r* = −0.31, *p <* .05), and MMSE score. However, these correlations were not significant when tested within groups (*p >* 0.05).

**Figure 11:**
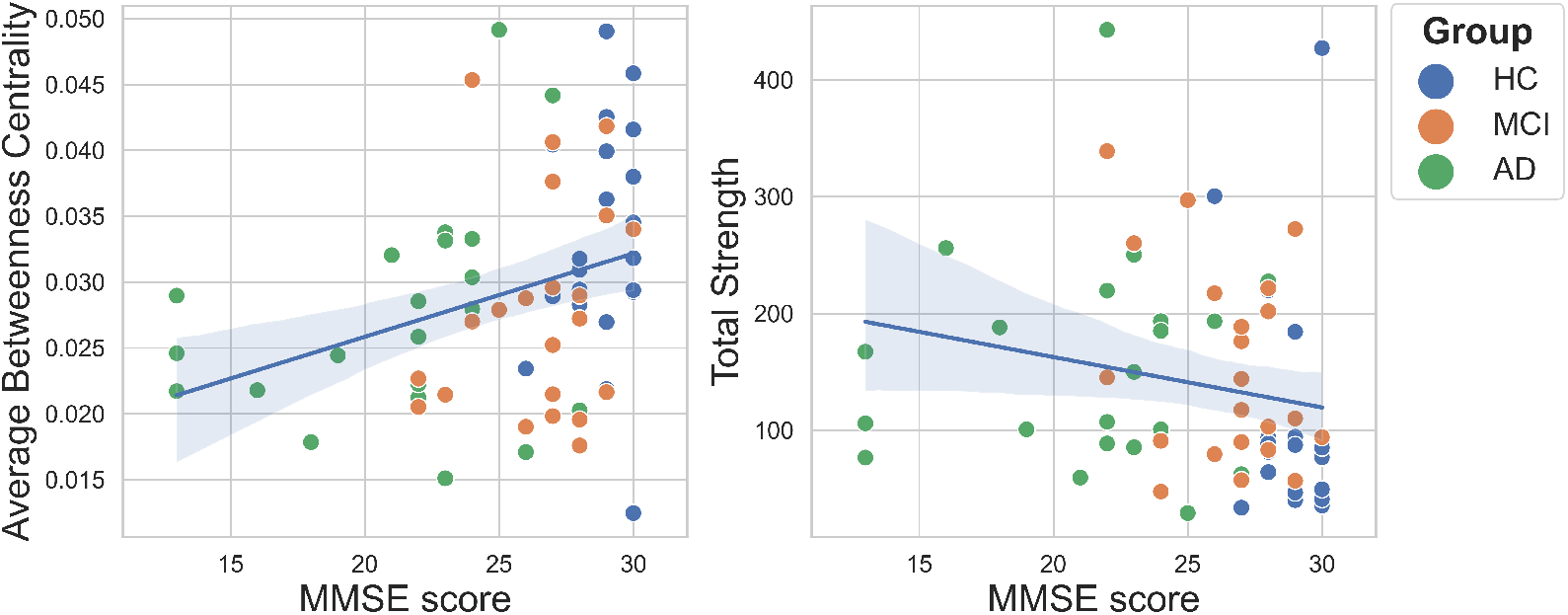
Correlations between average betweenness centrality and total strength, and Mini-Mental State Examination score (*p* values corrected using FDR).

## 4 Discussion

Dementia has been described as a ‘disconnection syndrome’, wherein large-scale disruptions in functional connectivity are clinically relevant: they are thought to emerge decades before symptom onset (Manno et al., 2019; Pereira, 2020), respond to medication (Lorenzi et al., 2011; Rizzi et al., 2022), are observed in mouse models (Zott et al., 2019), and mirror the spread of AD pathology (Griffa et al., 2013; Palmqvist et al., 2017). In this study, we applied a graph theory approach to quantify resting state functional connectivity, as measured by HD-DOT, in individuals with MCI, AD, and in age-matched healthy controls, which to our knowledge has not been previously explored. Such network-based approaches facilitate cross-study comparisons and provide insights into how dynamic properties emerge from underlying network topology. We identified group differences in network organisation at both global and local levels, alongside distinct profiles of key network hubs characterising each group. Contrary to our hypotheses, functional networks in both MCI and AD were hyperconnected, exhibiting greater density and strength relative to controls. While both hypo- and hyperconnectivity have been reported in the literature (Tijms et al., 2013), these alterations do not preclude the presence of reduced cerebral perfusion, which has been commonly observed in AD (Yeung & Chan, 2020). Interestingly, no differences were found between MCI and AD groups across networks metrics, suggesting that hyperconnectivity may emerge in the early stages of cognitive impairment. Both clinical groups were also associated with increased global efficiency, reflecting shorter path lengths and subsequently, more direct communication between brain regions. This finding is consistent with previous structural (Vermunt et al., 2020) and functional MRI (Agosta et al., 2011; Krienen & Buckner, 2009; Pievani et al., 2011; Supekar et al., 2008), as well as PET studies (Seo et al., 2013). The combination of shorter pathlengths, increased degree density, and higher total strength, suggests that networks in MCI and AD are more globally connected but in a disorganised fashion - a hallmark of disrupted metastability (Córdova-Palomera et al., 2017; Penalba-Sánchez et al., 2023). The further absence of consistent key network hubs in AD suggests a breakdown of specialised hierarchical organisation and an imbalance between local and global connectivity, which is somewhat preserved in MCI. Although we did not observe changes in clustering in the AD group, as has been reported with fMRI, a lack of network hubs in AD is a well-established finding (Tijms et al., 2013). Consistent with a loss of functional organisation, we also observed a positive correlation between MMSE score and average betweenness centrality, indicating that the number of nodes lying on the shortest paths between regions decreases with greater cognitive impairment. Taken together, these results suggest a shift toward more distributed network architectures in both AD and MCI, characterised by the recruitment of additional brain regions and a loss of functional specialisation.

Changes in connectivity can reflect both maladaptive and compensatory mechanisms, so the functional significance of the observed hyperconnectivity in MCI and AD is unclear. According to the dual-stage model of AD progression (Finn et al., 2015), early hyperconnectivity – driven by disease and gene-related factors – is thought to be compensatory, arising in response to declining MTL-PFC connectivity (Berron et al., 2020). In this model, the brain initially attempts to recruit alternative pathways to preserve cognitive function in the short term. Increased local connectivity supports communication near sites of damage, while increased global connectivity reflects the engagement of additional network hubs to maintain overall network function. At this stage, regions not yet burdened by tau pathology exhibit an imbalance in neuronal excitability, characterised by reduced inhibitory GABAergic activity and increased excitatory glutamatergic activity. This heightened neuronal excitability promotes tau accumulation and subsequently, amyloid deposition (Penalba-Sánchez et al., 2023), consistent with the ‘activity causes damage’ hypothesis (Bero et al., 2011). Amyloid pathology in turn impairs neurovascular coupling by reducing the production of vasodilatory factors, leading to hypoperfusion and hypoconnectivity in later stages of the disease. Thus while early hyperconnectivity may be initially adaptive, it can ultimately exacerbate disease progression by increasing vulnerability to tau and amyloid deposition (Bonanni et al., 2021). The increased PFC connectivity observed in the present study aligns with this framework, as functional alterations are thought to precede structural degeneration (Teipel et al., 2015) and frontal regions are among the last to accumulate tau and undergo atrophy (Braak & Braak, 1995). The relatively high MMSE scores of the AD group (*μ* = 21.3) and the lack of differences in network metrics between MCI and AD groups suggests that hyperconnectivity may indeed reflect an early adaptive mechanism, particularly as PFC atrophy is typically associated with lower MMSE scores (Salat et al., 2001). The observed loss of networks hubs for clustering and efficiency in AD, may indicate the onset of impaired global connectivity, as reported in later stages (Tijms et al., 2013), while local connectivity is preserved. Another possibility is that systemic physiological components contaminated the cortical signal in clinical groups. Prior work has demonstrated that even after regressing the extracerebral signals captured by short channels from the cortical signals captured by longer channels, residual systemic influences can still artificially inflate estimates of functional connectivity calculated using NIRS data (Abdalmalak et al., 2022). While there were no group differences between the percentage of good channels per channel distance, which excludes the possibility of a disproportionate contribution of short-channels in clinical groups, it remains possible that systemic physiology exerts a greater influence in these populations. For example, controlling for cardiovascular factors has been shown to eliminate age-related effects on resting state amplitude fluctuation measured using fMRI (Tsvetanov et al., 2020). Thus, it cannot be excluded that vascular alterations, rather than purely neural mechanisms, contributed to the present hyperconnectivity.

While there is no prior work using HD-DOT, previous studies using NIRS have indeed identified alterations in functional connectivity in dementia, though there is no clear consensus on the nature of these changes. This is in part owing to the limited number of studies published in the field and the heterogeneity in methodology across these studies (Albrecht et al., 2025; Bray et al., 2025). Findings are variable: both increased and decreased connectivity have been observed in MCI and AD (Butters et al., 2023). For example, some studies have reported reduced PFC connectivity in MCI (Ghafoor et al., 2019) and in AD (Keles et al., 2022) compared to controls, although these differences were not tested for significance. Additionally, reduced whole-brain connectivity (Zhang et al., 2022) and diminished effective connectivity between distal regions of the brain, e.g. PFC and the occipital lobe, have been reported in MCI (Bu et al., 2019). Conversely, *increased* inter-hemispheric PFC connectivity in MCI (Nguyen et al., 2019), and higher prefrontal spectral entropy in AD (Ferdinando et al., 2023) have also been reported. Some studies, however, find no differences in connectivity (G. Yang et al., 2025; Zheng et al., 2023). In the only study directly comparing MCI and AD, no differences in whole-brain functional connectivity were found (Niu et al., 2019), which aligns with the present findings. It is challenging to compare the present results with these studies for a number of reasons, which are detailed in Albrecht et al., 2025. Many studies used low-density systems with a single optode per region of interest (e.g. Bu et al., 2019). Whilst functional connectivity is generally more sensitive to diversity across regions than within the same region, the higher resolution in the present study allows for a more accurate representation of brain activity and a finer delineation of brain networks. Additionally, no prior studies used subject-specific anatomical priors for signal reconstruction which may have lead to the under-estimation of the haemodynamic signal. Moreover, whilst many studies measured functional connectivity using the average Pearson’s correlation coefficient between brain regions, graph theory offers a more comprehensive quantification of the *structural* organisation of the network. The only study to apply graph theory did so (Ghafoor et al., 2019) with a low-density system, perhaps lacking the spatial resolution necessary for detailed graph analysis.

Turning instead to the large body of work using fMRI, a general shift from early-stage hyperconnectivity, to later-stage hypoconnectivity in dementia has been documented (albeit with substantial individual variability; Dickerson et al., 2005; Pini et al., 2025). Looking specifically to within-PFC connectivity, evidence is mixed for MCI but AD is more consistently associated with hypoconnectivity (Ibrahim et al., 2021). It is uncertain whether connectivity measured by different modalities captures the same underlying construct (Tijms et al., 2013). The HD-DOT signal primarily reflects local haemodynamic activity within the microvasculature which occurs in superficial cortical layers, whereas fMRI captures generally broader activity, including that from deeper brain regions (Sasai et al., 2012). Therefore, while the present results may reflect locally increased PFC connectivity, they do not exclude global network disruption or disruption in deeper brain regions, as observed with fMRI (Li et al., 2014). This interpretation further aligns with the potential preservation of local connectivity in AD despite the onset of impaired global connectivity, as evidenced by a loss of network hubs. A key finding across fMRI studies is reduced PFC–MTL coupling and selective vulnerability of long-range connections (Berron et al., 2020). However, these connections cannot be explored using the current HD-DOT data due limited field-of-view and depth penetration, as the MTL is not imageable by NIRS. Whilst there is value in focusing on the PFC, as it is thought to be crucial in maintaining cognitive function during neurodegeneration (Jobson et al., 2021), doing so paints a limited picture of the complex interplay between the brain areas affected in dementia. The MTL atrophy rating scale used to rate hippocampal atrophy in the present study also may not serve as a relevant marker of structural change that can be directly compared with the functional changes observed in the PFC as the MTL and PFC differ in the timeline of their structural and functional changes (Jobson et al., 2021). Distinct alterations between AD and MCI have also been identified across several large-scale functional brain networks (Li et al., 2014). One network of particular interest, the DMN, has been well-characterised using fMRI. This network is typically more active during rest and deactivates during task states (Raichle et al., 2001). Altered connectivity within the DMN has been widely reported in AD (Ereira et al., 2024; Toussaint et al., 2014) and has shown strong potential for diagnosis (Ibrahim et al., 2021), as it is the first network affected by amyloid deposition (Buckner et al., 2009; Mintun et al., 2006). Although DMN connectivity was not captured in the present study, the medial PFC – part of the DMN – emerged as an important centrality hub in both MCI and AD groups and a greater number of regions in the DMN were important hubs in these groups compared to healthy controls. It has been proposed that key hubs in the DMN may be selectively vulnerable in AD, as signal fluctuations in the DMN have been found to negatively correlate with amyloid burden (Scheel et al., 2021). Furthermore, the majority of important hubs for AD and MCI were located in the left hemisphere, while in controls, these were located primarily in the right hemisphere. Such changes in frontal lateralisation have been observed in both NIRS and (Gjonaj et al., 2025) and fMRI (Liu et al., 2018), suggesting functional brain reorganisation in dementia.

There is a growing push to standardise dementia stratification using biologically-based diagnostic criteria. For example, the ATN framework for AD classifies individuals according to objective markers of amyloid, tau and neurodegeneration (Jack et al., 2016). The inclusion of a marker of brain *function* (Finn et al., 2015) into such classifications remains a topic of debate. To this end, functional connectivity may capture distinct clinical phenotypes present early in the disease course which are not captured by traditional markers such as structural atrophy (Pini et al., 2025). Yet incorporating such a functional marker would only be possible using tools which are accessible and scalable using current infrastructures. The present work demonstrates the potential of HD-DOT for this purpose. The majority of data collection took place in the home setting and we found no difference between the quality of the data collected in the home and that collected in a for-purpose clinical room at the University of Cambridge. The practical advantages of HD-DOT make it particularly well-suited for scalable clinical use, facilitating widespread deployment both in settings outside of the clinic and at multiple time points to support personalised treatment and diagnostic approaches. Nevertheless, analysis pipelines for HD-DOT remain largely unstandardised (Yücel et al., 2021) and current methods rely on computationally-expensive and inaccessible techniques like photogrammetry and MRI, which curtails its practicality. Efforts can be made to overcome these challenges though. Whilst assuming a standard brain size and shape likely introduces error (Srinivasan et al., 2023), as has been done in all previous work applying NIRS in dementia, acquiring a subject-specific MRI is often impractical. For such cases, an *atlas* for dementia could be developed, as has been done for infants (Collins-Jones et al., 2021). An appropriate head model could be chosen based on various parameters such as head size, age, and symptom history.

## Limitations

The present study has certain limitations. Firstly, we used what is considered to be a relatively short resting state duration. Typically, the optimal acquisition time for resting state fMRI is around ten minutes, as test-retest reliability of functional connectivity measures improves with longer scan durations (Birn et al., 2013). However, this is likely because fMRI has a lower sampling rate of around 0.5-1 Hz, compared to the generally higher sampling rates of NIRS and HD-DOT, and that of 12.5 Hz used in the present study. In fact, previous work using NIRS has shown that as little as 30 seconds of resting state data can distinguish between MCI and AD (D. Yang & Hong, 2021). Secondly, as the HD-DOT signal is a multiplexed combination of neural, haemodynamic and noise components, disentangling vascular and neural contributions to the signal is not possible (Ereira et al., 2024). This is particularly pertinent to dementia where we expect underlying disease-related vascular changes to influence cerebral haemodynamics (de la Torre, 2012). The degree to which the presently-observed differences in functional connectivity are due to such vascular changes or reflect true neural activity is thus unknown. This issue is not unique to HD-DOT however, it is also inherent to fMRI (Drew, 2019), and can be addressed by the use of simultaneous EEG-NIRS paradigms.

## Future work

To our knowledge, this is the first study that has applied HD-DOT to dementia, opening up several avenues for future research in the area. Dementia leads to widespread dysfunction across many cortical and sub-cortical brain regions (Raji et al., 2009). Evaluating whole-brain functional connectivity would therefore help to provide a more comprehensive understanding of the patterns observed in the present work. This could be achieved using newly-developed whole-head HD-DOT systems (Collins-Jones et al., 2024). In addition, using task-based data instead of the resting state may tease out more pronounced differences between groups and reveal more subtle within-group variations (Nguyen et al., 2019). The ONAC study employed a range of task paradigms, the data of which remain to be explored, and also used broadband NIRS to assess neurometabolism (Acharya et al., 2025). Finally, alternative computational methods could be applied to our data to provide further insights into networks beyond their structural organisation. For example, computing effective connectivity would determine the directionality of functional relationships between brain regions (Ereira et al., 2024). A more mechanistic approach would be dynamic causal modelling (Tak et al., 2015), which explicitly models the processes underlying these relationships. This type of modelling has not yet been applied to HD-DOT as far as we are aware.

## 5 Conclusion

In summary, we have shown that HD-DOT is capable of detecting differences in both local and global functional connectivity in the resting state between individuals with AD and MCI, and healthy controls.

These findings suggest increased network density and strength in AD and MCI, alongside a loss of hierarchical organisation and key network hubs as the disease progresses. Through this work, we have also demonstrated the feasibility of using HD-DOT for high-quality, at-home assessments of brain function. Future work should employ longitudinal study designs to assess the sensitivity of HD-DOT to disease-related changes, and should prioritise the development of standardised analysis and preprocessing pipelines to improve the useability of HD-DOT in clinical settings.

## 6 Data and Code Availability

The code is available at http://www.github.com/emiliavioletb/ImagingNeuroscience. Data are available upon reasonable request from accredited researchers, in accordance with our ethical approval.

## 7 Author Contributions

Emilia Butters: Conceptualisation, Methodology, Software, Formal analysis, Investigation, Data curation, Writing—original draft, Writing—review & editing, Visualisation, Project administration, and Funding acquisition. Liam Collins-Jones: Methodology, Software, Writing—review & editing. Rickson Mesquita: Methodology, Software, Formal analysis, Writing—review & editing, Visualisation. Deepshihka Acharya: Software, Writing—review & editing. Elizabeth McKiernan: Investigation, Resources, Writing—review & editing. Axel Laurell: Investigation, Writing—review & editing. Audrey Low: Investigation, Writing—review & editing. Sruthi Srinivasan: Validation. John O’Brien: Conceptualization, Resources, Writing - Review & Editing, Supervision, Funding acquisition. Li Su: Conceptualization, Resources, Writing - Review & Editing, Supervision. Gemma Bale: Conceptualisation, Writing—review & editing, Supervision, Funding acquisition.

## 10 Acknowledgments

We would like to thank all the subjects and informants who took part in this study, and RHM for kindly lending his car for data collection.

## 11 Appendix

### 11.1 Data quality between testing locations

**Figure A1:**
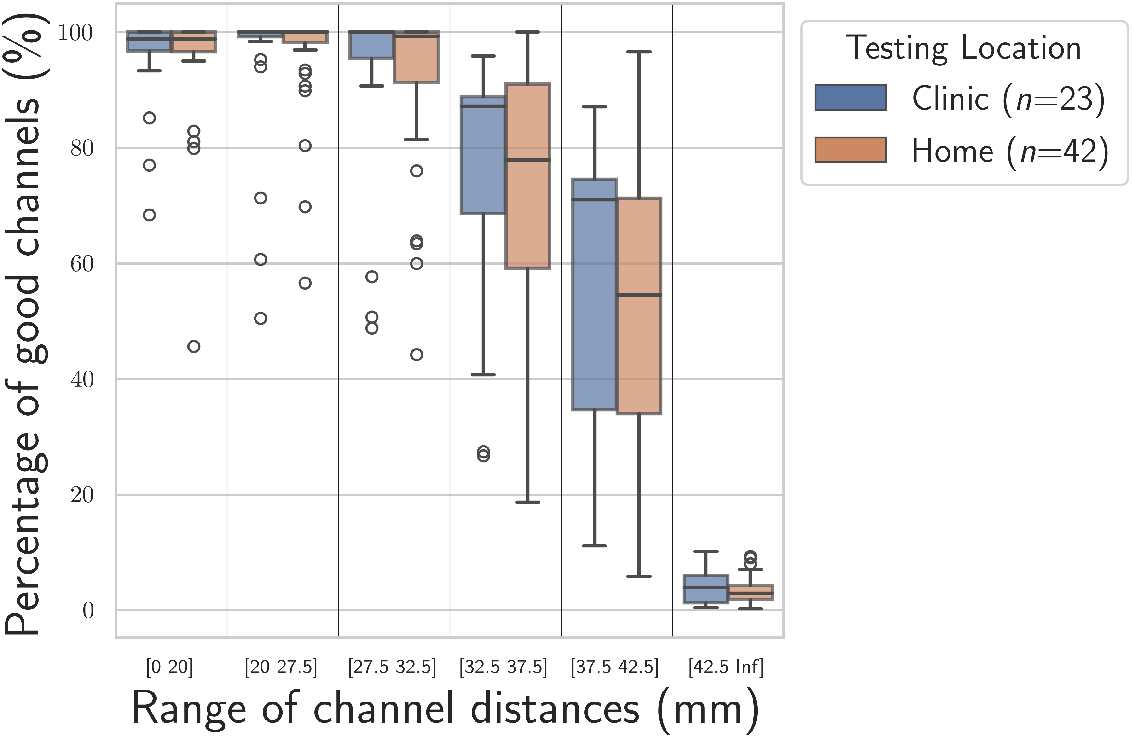
No significant differences in the percentage of good channels across channel distances between testing locations.

**Table A1:**
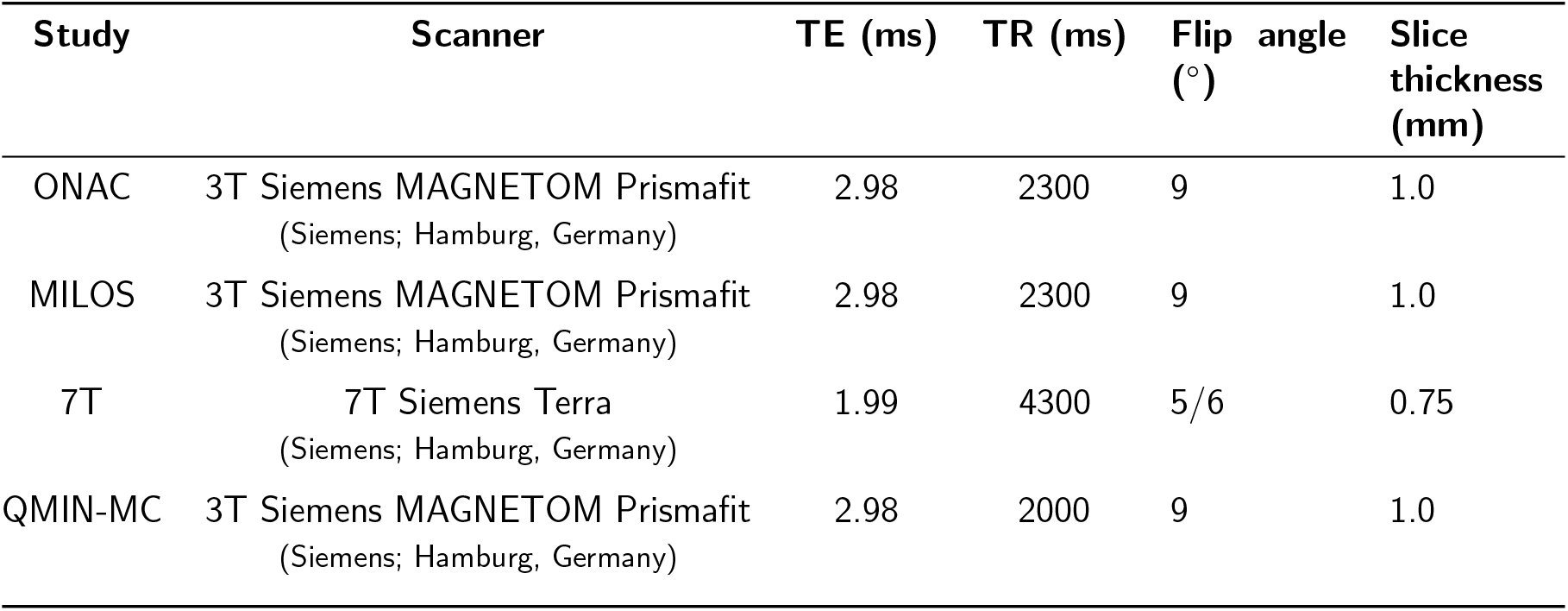
MRI sequence parameters across studies.

### 11.2 Details of MRI acquisition

Across the MCI and AD groups, a total of 43 participants required MRI scans for the creation of head models. Of these, 26 were scanned through the ‘Optical Neuroimaging and Cognition’ study, four through the ‘MultiModal Imaging in Lewy-Body Disorders’ study, four through the ‘7T MRI for Dementia with Lewy Bodies’, and nine through the ‘Quantitative MRI in the NHS Memory Clinics’ study. The respective sequence parameters are detailed in Table A1.

Scans acquired at 7T were de-noised using a custom script (www.github.com/benoitberanger/mp2rage) and bias field corrected using SPM12 (www.fil.ion.ucl.ac.uk/spm) with light regularisation (0.001).

### 11.3 Optode array coverage

There were no differences in the number of sensitive parcels across groups (*H*(2) = 2.10, *p* = 0.35), as shown in Figure A2a.

**Figure A2:**
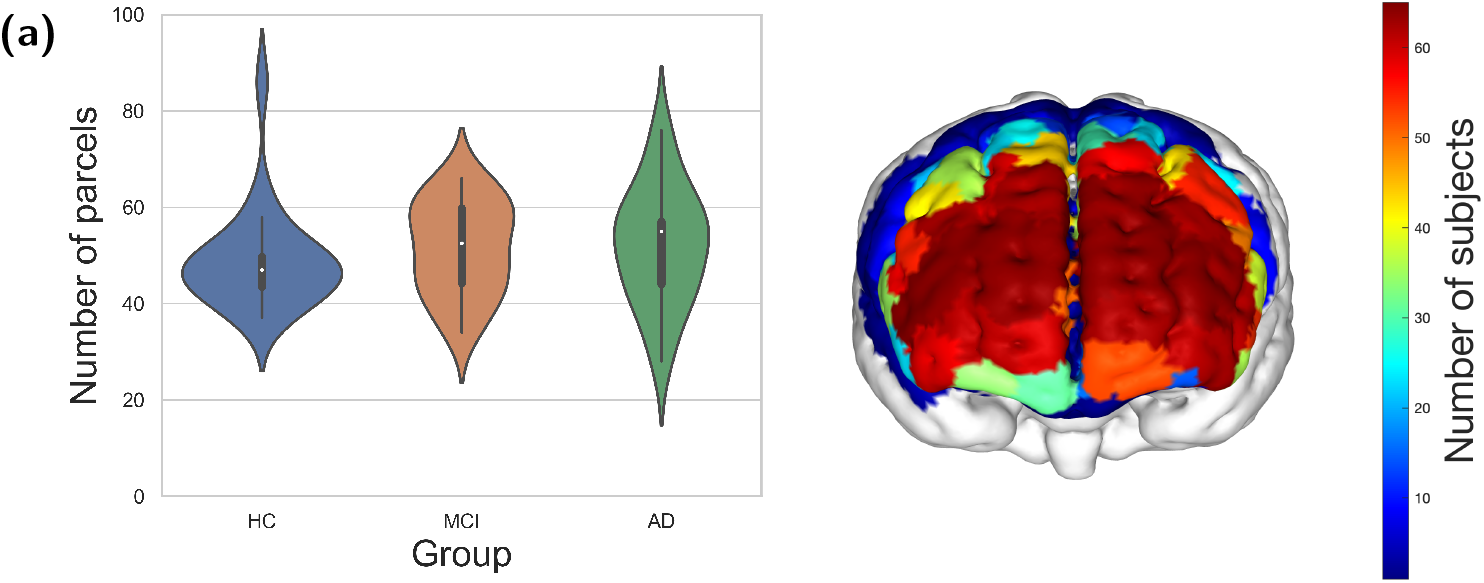
Coverage of the frontal array. (a) The number of sensitive parcels for each subject per group. (b) Histogram of the number of subjects sensitive to all frontal parcels.

