## Supplementary Table 1 for "Brain Network Analysis in Alzheimer’s Disease and Mild Cognitive Impairment using High-Density Diffuse Optical Tomography"

### Supplementary material

Table S1: Summary of neuropsychological test scores across study groups.

| | **HC** (*n*=22) | **AD** (*n*=21) | **MCI** (*n*=22) | **p** |
|---|---|---|---|---|
| *Attention/processing speed* | | | | |
| Digit span | 9.77 ± 2.04 | 5.43 ± 2.50 | 8.19 ± 1.74 | **0.00**[a,b,c] |
| TMT-A time (s) | 41.6 ± 12.9 | 117 ± 87.0 | 50.8 ± 21.3 | **0.01**[b,c] |
| TMT-A errors | 0.09 ± 0.29 | 0.33 ± 1.08 | 0.20 ± 0.6 | 0.76 |
| *Executive function* | | | | |
| Interference sensitivity | 2.91 ± 0.29 | 1.81 ± 1.26 | 2.95 ± 0.21 | **0.00**[b,c] |
| Inhibitory control | 2.64 ± 0.71 | 1.29 ± 1.12 | 2.38 ± 1.05 | **0.01**[b,c] |
| TMT-B time (s) | 85.1 ± 41.6 | 54.2 ± 93.7 | 114 ± 46.2 | **0.02**[b,c] |
| TMT-B errors | 15.3 ± 2.22 | 1.19 ± 2.59 | 2.00 ± 2.60 | 0.45 |
| *Motor function* | | | | |
| UPDRS Part III | 3.14 ± 3.11 | 7.67 ± 7.14 | 4.82 ± 3.95 | 0.37 |
| *Olfactory function* | | | | |
| B-SIT | 9.09 ± 2.64 | 6.10 ± 2.79 | 7.65 ± 2.89 | 0.26 |
| *Colour discrimination* | | | | |
| Farnsworth total error | 8.63 ± 18.2 | 20.2 ± 21.4 | 6.84 ± 9.54 | **0.04**[c] |
| *Visual hallucinations* | | | | |
| Pareidolia score | 37.7 ± 8.44 | 34.7 ± 8.87 | 39.5 ± 0.84 | **0.01**[b,c] |
| Presence of hallucinations (%) | 0 | 9.50 | 9.10 | 0.98 |
| *Anxiety & depression* | | | | |
| HADS | 5.18 ± 4.53 | 7.33 ± 3.63 | 9.58 ± 4.50 | 0.37 |
| GDS | 1.50 ± 2.79 | 2.43 ± 2.13 | 3.64 ± 3.38 | 0.37 |
| *Informant questionnaires* | | | | |
| BADLS | n/a | 9.95 ± 7.02 | 2.50 ± 3.61 | **0.00** |
| CDR | n/a | 6.91 ± 3.43 | 4.06 ± 2.72 | **0.01** |
| CBI | n/a | 42.0 ± 17.6 | 26.6 ± 14.0 | **0.01** |
| CAF | n/a | 1.19 ± 2.56 | 3.50 ± 1.66 | 0.50 |
| CAF (One day) | n/a | 2.48 ± 2.84 | 0.55 ± 1.24 | **0.01** |
| DCFS | n/a | 9.71 ± 3.12 | 9.31 ± 2.20 | 0.37 |
| NPI | n/a | 18.2 ± 6.74 | 9.82 ± 10.3 | 0.07 |

Shown as mean ± standard deviation or percentage (%).
P-value indicates overall group-level comparison. Superscript letters denote significant pairwise group comparisons: [a] HC vs MCI, [b] HC vs AD, [c] MCI vs AD.
MMSE, Mini-Mental State Examination (Folstein et al., 1975); MoCA, Montreal Cognitive Assessment (Nasreddine et al., 2005); TMT, Trail Making Test (Partington & Leiter, 1949); UPDRS, Unified Parkinson's Disease Rating Scale (Goetz et al., 2008); B-SIT, Brief Smell Identification Test (Doty et al., 1996); Farnsworth; D-15 color arrangement test. Pareidolia; the Noise Pareidolia test (Mamiya et al., 2016); HADS, Hospital Anxiety and Depression scale (Zigmond & Snaith, 1983); GDS, Geriatric Depression Scale (Yesavage & Sheikh, 1986); BADLS, Bristol Activities of Daily Living Scale (Bucks et al., 1996); CDR, Clinical Dementia Rating (Morris, 1993); CBI, Cambridge Behavioural Inventory (Wear et al., 2008); CAF, Clinician Assessment of Fluctuation (Walker et al., 2000); DCFS, Dementia Cognitive Fluctuation Scale (Lee et al., 2014). NPI, Neuropsychiatric inventory (Cummings et al., 1994).
